# Particulate Matter emission sources and meteorological parameters combine to shape the airborne microbiome communities in the Ligurian coast, Italy

**DOI:** 10.1101/2020.08.06.239947

**Authors:** Giorgia Palladino, Pietro Morozzi, Elena Biagi, Erika Brattich, Silvia Turroni, Simone Rampelli, Laura Tositti, Marco Candela

## Abstract

Here we explore how the chemical composition of particulate matter (PM) and meteorological conditions combine in shaping the air microbiome in a heavily inhabited industrial urban settlement. During the observation time, the air microbiome was highly dynamic, fluctuating between different compositional states, likely resulting from the aerosolization of different microbiomes emission sources. This dynamic process depends on the combination of local meteorological parameters and particle emission sources, which may affect the prevalent aerosolized microbiomes. In particular, we showed that, in the investigated area, industrial emissions and winds blowing from the inlands combine with an airborne microbiome that includes faecal microbiomes components, suggesting multiple citizens’ exposure to both chemicals and microorganisms of faecal origin, as related to landscape exploitation and population density. In conclusion, our findings support the need to include monitoring of the air microbiome compositional structure as a relevant factor for the final assessment of local air quality.

## Introduction

Airborne Particulate Matter (APM) is a complex system of particles in suspension in the atmosphere, usually defined as aerosol. Atmospheric aerosol is contributed by a multiplicity of sources of both natural and anthropogenic origin, including both biogenic and abiotic chemical components, and producing extremely complex and variable matrices that can be processed and solved for their origin using appropriate analytical processing and computational tools [1, 2]. In particular, the aerosol composition consists of a series of macrocomponents, which make up the mass of APM, as well as an even larger series of different trace components, the latter being of primary relevance as including the most toxic species and providing the highest chemical fingerprinting potential [3]. These aerosol bulk components can be emitted directly into the atmosphere (Primary Aerosol) or, otherwise, they can be abundantly produced within the atmosphere, following chemical reactions on gaseous precursors previously emitted (Secondary Aerosol). Primary Biological Aerosol (PBA), in short bioaerosol, represents the APM fraction including atmospheric particles released from the biosphere to the atmosphere [4]. PBA comprise living and dead organisms, their dispersal units (e.g. pollen and spores) as well as tissue fragments from decay processes [5, 6]. The overall mass contribution of PBA to conventional APM metrics is to date a very challenging task though some authors have recently estimated that it may account for about 16% of PM_10_ in different cities examined [7]. The PBA fraction including microorganisms is defined as “airborne microbiome” (AM) and represents a highly dynamic and diversified assemblage of active and inactive microorganisms [8]. Indeed, AM can originate from multiple terrestrial and marine sources – including resident microbiomes in soil, waterbodies, plants and animal faeces [4, 9] – whose relative importance depends season, location, altitude and meteorological and atmospheric factors. Further, in agricultural and suburban locations, other sources relevant to AM emissions are represented by man-made systems, such as agricultural waste, composting, and wastewater treatment plants. AM emission mechanisms include erosion or abrasive dislodgement from terrestrial sources and, from open waters, bubble-bursting at the air-water interface [10, 11]. PBA size spans from a few nanometres up to about a tenth of a millimetre [5], with bacteria-containing particles ranging around 2-4 μm in diameter [12] and accounting for “5–50% of the total number of atmospheric particles >0.2 μm in diameter” [13]. Due to the small size, AM can be transported over large distances, across continents and oceans, and reach the upper troposphere, where it actively contributes to ice nucleation and cloud processing [14]. In the troposphere, the AM concentration ranges from 10^2^ to 10^5^ cells/m^3^ [12], being the densest in the planetary boundary layer, whose thickness depends on micrometeorological factors and geographic location, with marked daily and seasonal fluctuations [15, 16]. In particular, the near-ground AM is the one most influenced by local sources, including local meteorology and atmospheric composition. AM is then removed from the troposphere by wet and dry deposition processes. The former is the major sink for atmospheric aerosol particles, in the form of precipitation [17], while the latter, being less important on the global scale, is particularly relevant with respect to local air quality [4, 8].

Recently, an increasing perception of the strategic importance of PBA – AM in particular – for the Earth system and, ultimately, for the planet and human health, has arisen [18–20]. For instance, besides its relevance to atmospheric processes, AM has been found to control the spread of microorganisms over the planet surface, affecting the geographical biome, with key implications on agriculture and, ultimately, human health. This awareness raised concern about the potential impact of anthropic activities on PBA and, in particular, on the AM fraction. For example, changes in aerosol composition due to extensive human influence on the planetary scale give rise to air pollution, the inherent modification of atmospheric reactivity and, ultimately, climate change [21]. These factors may likely interfere with AM, shaping its structure and dispersion throughout the troposphere, with direct consequences on the terrestrial biome [22]. However, as far as we know, the current state of knowledge on the connections between AM, atmospheric processes and atmospheric pollution is still fragmentary, especially due to the lack of a cross-cut approach. Therefore, in this work, we explore the ability of an interdisciplinary approach combining chemical speciation and metagenomics in shedding light on the complex relationships among abiotic and microbiome components of local ambient aerosol. The study is based on a series of about one hundred PM_10_ samples from a coastal district in north-western Italy, collected daily over six months, from February to July 2012, to cover the cold-to-warm seasonal transition. The chemical composition of each sample was obtained, and a receptor modelling approach was used to identify and quantitatively apportion the chemical species determined in the samples to their sources. Owing to the cutoff adopted in APM sampling, the samples were deemed suitable for total DNA extraction and microbiome characterization by Next-Generation Sequencing using the 16S rRNA gene as target. In our work, we were able to finely reconstruct the overall aerosol behaviour in an area affected by both natural and anthropogenic emission sources, determining the local bacterial microbiome from PBA contained in PM_10_ and its main features as a function of local meteorological and environmental characteristics.

## Materials and methods

### Site description

The PM_10_ samples treated in this work were collected in Savona, one of the main towns in the Ligurian region (**Figure 1**). The whole region overlooks the Tyrrhenian sea but is entirely occupied by the Appenninic range down to the coast, where only a narrow strip of plain is present. Therefore, the coastal area is densely inhabited and crossed by an extremely busy traffic road mainly connecting Italy to France. Besides being occupied by a medium-size heavily inhabited urban settlement, the Savona district also hosts a wide industrial area, including a coal-fired power plant active at the time of our experimental field activity and a large and very busy harbour. The climate of this site is classified as warm-temperate (Csa, according to Köppen and Geiger classification) [23–24] with an average annual temperature of 14.6°C and average precipitation of 910 mm (https://en.climate-data.org, accessed 28/07/2020). Intense northern winds characterize the circulation in winter [25], while sea-land breeze regimes prevail in the warm season, usually starting from March [26, 27].

**Figure 1.**
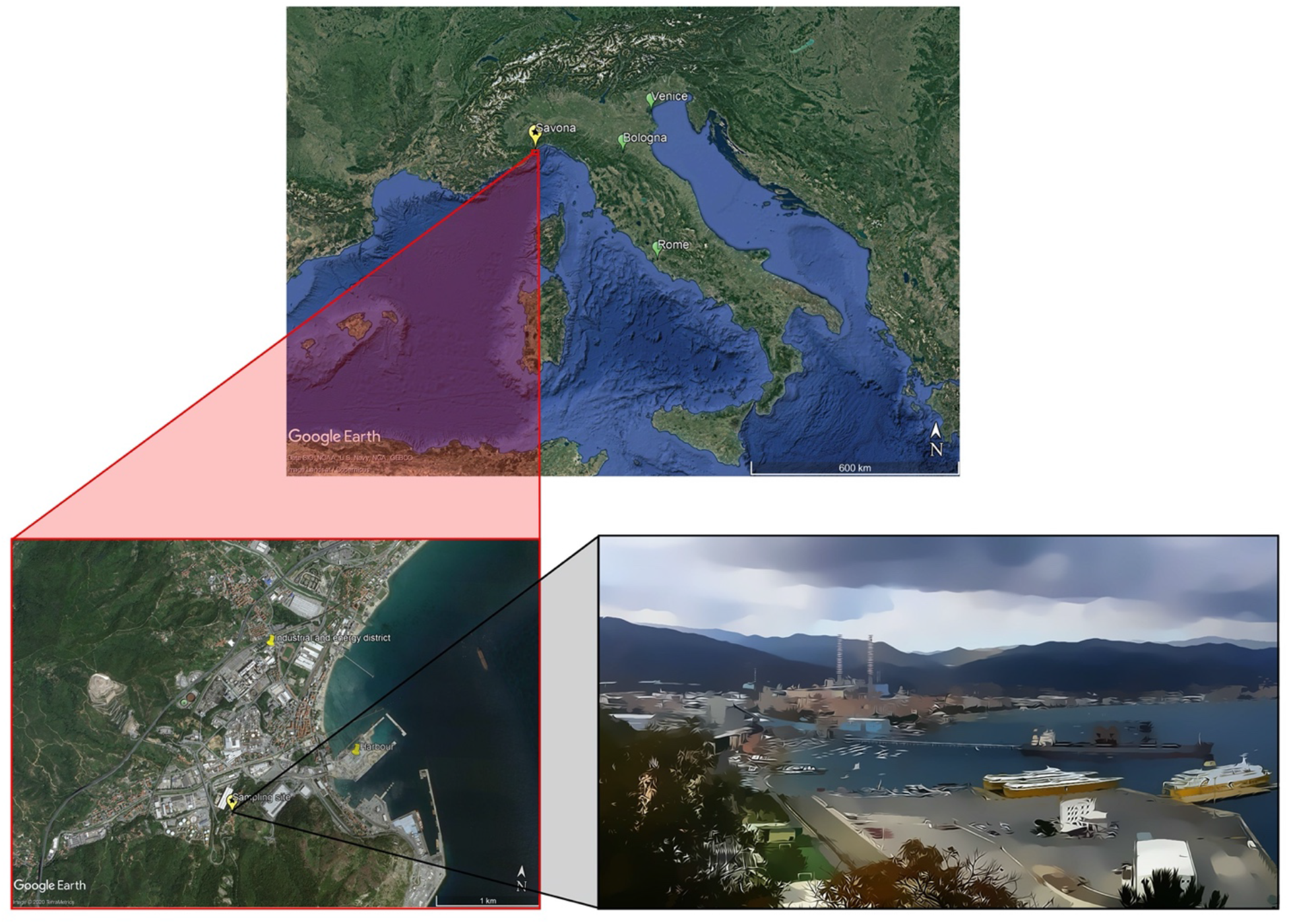
Location of the sampling site. Map providing the location of Savona in Italy (indicated with a yellow balloon with a star; other Italian cities are indicated with a green balloon) (top panel), and snapshot of the PM_10_ sampling site with a 3D view of the surroundings (bottom panel) (source: Google Earth; map data: SIO, NOAA, U.S. Navy, NGA, GEBCO, TerraMetrics).

### Sample collection and atmospheric parameters

A total of 184 daily PM_10_ samples were collected from February 1, 2012, until July 20, 2012 with low-volume samplers (SWAM Dual Channel, 55.6 m^3^ /day, FAI, Italy) to allow simultaneous collection of both quartz (Whatman ®QM-A quartz) and PTFE membranes (Whatman PM_2.5_ PTFE). Samples were stored frozen in the dark at −10°C until processing. In this work, PTFE membranes were used for gravimetry, ion chromatography and elemental analysis with particle induced X-ray emission and inductively coupled plasma mass spectrometry, while quartz membranes were used for the analysis of carbonaceous macrocomponents and microbiology. A subset of 98 samples, uniformly distributed across the sampling period, was used for the analyses reported in the present paper. During the sampling campaign, meteorological parameters were measured simultaneously on site using a Davis Vantage Pro2 Weather Station (Davis Instruments, Hayward, CA), placed in proximity of the PM_10_ sampler, for the measurement of temperature, pressure, relative humidity, rainfall, and wind direction and speed with a time resolution of 30 min. Subsequently, the data obtained were averaged on a daily scale (**Supplementary Table S1**), i.e. at the same time resolution as the PM_10_ samples, using the “openair” package [28] of the R software (version 3.6.1; https://www.r-project.org/).

### Chemical characterisation of the samples

Chemical characterization of PM_10_ filters was carried out using several analytical techniques. First, PM_10_ mass load (μg/m^3^) was determined by gravimetric analysis. Elemental and organic carbon were determined on quartz membranes by thermal-optical transmittance analysis (TOT), as previously described [29]. For inorganic speciation, several analytical techniques were performed on PTFE filter portions: Ion Chromatography (IC) for the determination of the main water-soluble ion composition (NH_4_^+^, K^+^, Mg^2+^, NO_3_^−^, SO4^2−^, Na^+^, Cl^−^, Ca^2+^, and a few low-level organic compounds, *i.e.* oxalates and methanesulfonate), and Particle Induced X-ray Emission (PIXE) and Inductively Coupled Plasma Mass Spectrometry (ICP-MS) for the simultaneous analysis of a series of metals and metalloids (Na, Al, Si, Cl, Ca, Ti, V, Cr, Mn, Fe, Ni, Cu, Zn, Br, Pb, Li, Co, Rb, Sr, Cd, La, Ce, Sb, Cs, Ba, Ti, Bi, As, Se, Sn). Elemental analysis by PIXE was carried out at the Tandetron 3 MeV of LABEC-INFN, Florence (Italy), according to the method previously reported [30]. Elemental analysis by ICP-MS was carried out according to the UNI EN 14902, 2005 for PM_10_ as an extension of the DL 155, 2010 in agreement with the EU Directive 2008/50/EC on ambient air quality and cleaner air for Europe. In order to prevent data redundancy, insoluble magnesium (Mg ins) and insoluble potassium (K ins) were calculated as the difference between PIXE and IC concentrations and replaced the corresponding elementary concentration data.

### Positive Matrix Factorization analysis

Positive Matrix Factorization (PMF) is an advanced multivariate factor analysis technique widely used in receptor modelling for the chemometric evaluation and modelling of environmental datasets [3, 31–36]. PMF allows the identification and quantification of the emissive profile of a receptor site, the monitoring site where an air quality station is operated. We applied EPA PMF 5.0 software [37]. The dataset was checked and re-arranged prior to PMF modelling according to the model criteria previously described [37] and, after data pre-processing, a concentration matrix of 98 samples × 25 variables was obtained. After careful evaluation of the input data and uncertainty matrices, an optimum number of factors was found by analysing the values of Q, a parameter estimating the goodness of the fit performed [38], and the distribution of residuals. In order to assess the reliability of the model reconstruction, measured (input data) and reconstructed (modeled) values together with the distribution of residuals were compared. Our results indicated a good general performance of the model in reconstructing PM_10_ (coefficient of determination equal to 0.79) for most variables. In order to confirm the results of receptor modelling, the origin of the air masses associated with the factors obtained was investigated through the creation of wind polar plots using the source contribution of the factors produced by PMF. In particular, polar plots were produced for each single PMF factor using the “openair” package of R [28], utilizing the conditional probability function (CPF) [39] with an arbitrary threshold set to the 75th percentile.

### Microbial DNA extraction, 16S rRNA gene amplification and sequencing

Microbial DNA extraction was performed on quartz membrane filter using the DNeasy PowerSoil Kit (Qiagen, Hilden, Germany) with the following modifications: the homogenization was performed with a FastPrep instrument (MP Biomedicals, Irvine, CA) at 5.5 movements per sec for 1 min, and the elution step was preceded by a 5-min incubation at 4°C [40, 41]. Extracted DNA samples were quantified with NanoDrop ND-1000 (NanoDrop Technologies, Wilmington, DE) and stored at −20°C until further processing. The V3-V4 hypervariable region of the 16S rRNA gene was PCR amplified in a 50-μL final volume containing 25 ng of microbial DNA, 2X KAPA HiFi HotStart ReadyMix (Roche, Basel, Switzerland), and 200 nmol/L of 341F and 785R primers carrying Illumina overhang adapter sequences. The thermal cycle was performed as already described [42], using 30 amplification cycles. PCR products were purified using Agencourt AMPure XP magnetic beads (Beckman Coulter, Brea, CA). Indexed libraries were prepared by limited-cycle PCR with Nextera technology and cleaned-up as described above. Libraries were normalized to 1 nM and pooled. The sample pool was denatured with 0.2 N NaOH and diluted to 6 pM with a 20% PhiX control. Sequencing was performed on an Illumina MiSeq platform using a 2 × 250 bp paired-end protocol, according to the manufacturer’s instructions (Illumina, San Diego, CA). Sequence reads were deposited in the National Center for Biotechnology Information Sequence Read Archive (NCBI SRA; BioProject ID XXXX).

### Bioinformatics and statistics

A pipeline combining PANDAseq [43] and QIIME 2 [44] was used to process raw sequences. DADA2 [45] was used to bin high-quality reads (min/max length = 350/550 bp) into amplicon sequence variants (ASVs). After taxonomy assignment using the VSEARCH algorithm [46] and the SILVA database (December 2017 release) [47], the sequences assigned to eukaryotes (*i.e.* chloroplasts and mitochondria) or unassigned were discarded. Three different metrics were used to evaluate alpha diversity – Faith’s Phylogenetic Diversity (PD whole tree) [48], Chao1 index for microbial richness, and number of observed ASVs – and unweighted UniFrac distance was used for Principal Coordinates Analysis (PCoA). Permutation test with pseudo-F ratio (function “adonis” in the “vegan” package of R), Kruskal-Wallis test or Wilcoxon rank-sum test were used to assess data separation. Kendall correlation test was used to determine associations between the PCoA coordinates of each sample and the factors identified by the PMF analysis. P-values were corrected for multiple testing with the Benjamini-Hochberg method, with a false discovery rate (FDR) ≤ 0.05 considered statistically significant. All statistical analyses were performed in R using “Made4” [49] and “vegan” (https://cran.r-project.org/web/packages/vegan/index.html) packages. Clustering analysis of family-level AM profiles, filtered for family subject prevalence of at least 20%, based on the SILVA taxonomy assignment, was carried out using hierarchical Ward-linkage clustering based on the Spearman correlation coefficients. We verified that each cluster showed significant correlations between samples within the group (multiple testing using the Benjamini–Hochberg method) and that the clusters were statistically significantly different from each other (permutational MANOVA using the Spearman distance matrix as input, function adonis of the vegan package in R).

Additionally, PANDAseq assembled paired-end reads were also processed with the QIIME1 [50] pipeline for OTUs (Operational Taxonomic Units) clustering based on 97% similarity threshold. Taxonomy was then assigned using the SILVA database. OTUs were processed through the R package “MaAsLin2” [51] to determine their association with microbial clusters. Kruskal-Wallis test was used to find OTUs whose relative abundance was significantly different among microbial clusters. The resulting OTUs were taxonomically assigned against the NCBI 16S rRNA database using the BLAST algorithm (https://blast.ncbi.nlm.nih.gov/).

## Results

### Particulate Matter emission sources and atmospheric parameters

The PMF model application on PM_10_ samples resulted in a solution with an optimum number of seven source factors at the receptor site, *i.e.* the station where the PM_10_ samples were collected. Like other multivariate methods, these factors correspond to linear combinations of the original compositional parameters, each potentially identifiable as an emission source profile. The fractional contribution per sample for each of the seven factors is reported in **Supplementary Table S2**.

In order to associate the factors with specific emission sources, prior knowledge about the receptor site (Savona, Italy) was used together with a critical analysis of the factor fingerprints (**Figure 2A**). Moreover, the percentage contribution of the seven identified sources to the total variable was reported (PM_10_, **Figure 2B**). As a result, the seven factors extracted by PMF analysis can be described as follows. Factor 1 is characterized by the prevalence of elements attributable to the geochemical composition because of the high percentages of Si, Al, and Ti. Therefore, this factor was identified as “crustal material and road dust resuspension”, deriving from the soil and/or road surface [34, 52]. Factor 2 is linked to organic carbon (OC), Cu, Zn, Cr, and K^+^. OC and K^+^ are strictly related to combustion processes, including biomass burning, as previously described [53]. Cu, Zn, and Cr are associated with traffic: Cu and Cr are well-known tracers of the brakes of motor vehicles, while Zn is known as a tracer of tire wear [54–56]. Therefore, this factor was identified as a combination of “traffic and biomass burning” sources. Factor 3 is mainly associated with NO_3_^−^ from gas-to-particle conversion of NO_x_ (g) in the atmosphere to which traffic and other high-temperature combustion processes may contribute [57, 58]; as such it can hardly be attributed to a single well-defined source, especially in such a complex emissive scenario. Therefore, this factor was identified collectively as “secondary nitrate”. Factor 4 relates to SO_4_^2−^ and NH_4_^+^ from gas-to particle reactions, leading to secondary ammonium sulphate [59–61]. Similarly to secondary nitrate, this component can be contributed by various sources (both natural and anthropogenic) due to the multiplicity of fossil fuel sources of the precursor gaseous SO_2_ and the ubiquity of NH_3_ (g) [62, 63]. Therefore, this factor was collectively identified as “secondary ammonium sulphate”. Factor 5 is associated with Na, Mg ins, V, and Ni. The distinctive association of V and Ni reveals emissions attributable to the combustion of heavy oil [64–66]. The association of these species with Na and Mg suggests a “naval-maritime transport” source. Factor 6 is mainly characterized by high scores of Pb, K ins, Zn, OC, and elemental carbon (EC). The fine particles produced by coal combustion are characterized by significant fractions of OC and K together with typical elements such as Zn, while other semi-volatile elements condense on the surface of fine particles of K ins [67]. Therefore, this factor was identified as “coal burning”. Factor 7 is connected to a large score of Cl^−^, Na, and Mg^++^, and clearly identified as “sea spray” aerosol [68].

**Figure 2.**
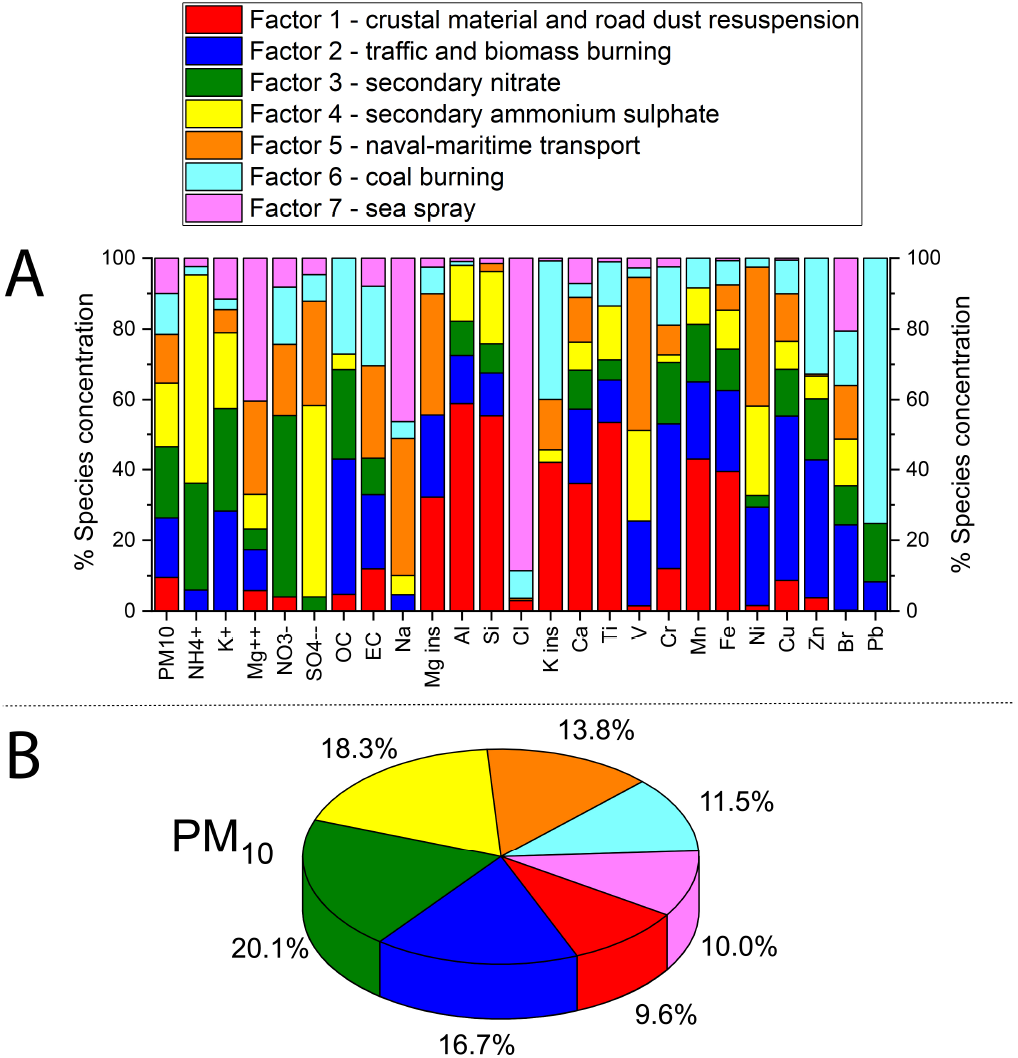
Emission sources identified by PMF analysis. (**A**) Stacked bar chart of the percentage concentration of each chemical species contributing to each of the seven factors that represent the chemical profile of each source identified in the PMF model. (**B**) Pie chart representing the contribution of the seven sources to PM10 mass. The seven factors were identified as reported at the top.

In order to confirm the PMF analysis results, the origin of the polluted air masses was investigated by analyzing the PMF factors as a function of wind direction, calculating the respective cumulative distribution functions and generating the corresponding wind polar plots. This method associates the emissive profile obtained by PMF with wind direction and intensity to which the receptor site is downwind. The plots obtained are shown in **Supplementary Figure S1**. In particular, factors 1, 3, 4, and 6 (respectively crustal material and road dust resuspension, secondary nitrate, secondary ammonium sulphate, and coal burning) are associated with winds blowing from the inland towards the coast covering traffic and industrial sources. Factor 5 (naval-maritime transport) is oriented downwind from the sea, confirming that it is associated with the fuel oil used for sea shipping. Finally, while factor 2 shows a local origin indicating sources in the proximity of the receptor site, factor 7 is meridionally oriented, indicating once more a marine origin. It should be noted, however, that, unlike factor 5 characterized by elements typical of the submicron fraction likely flushed back and forth by sea-land breezes from the harbor, factor 7 is associated with coarse particles requiring different meteorological conditions (possibly more intense winds from the open sea in order to sustain heavier particles).

### AM overall composition

Next generation sequencing of the V3-V4 hypervariable region of the 16S rRNA gene from the total microbial DNA extracted from PM_10_ air filters resulted in 98 samples containing more than 1 000 reads per samples which were retained for the rest of the study, for a total of 797 781 high-quality sequences with an average of 8 058 ± 3 410 (mean ± SD) paired-end reads per sample, binned into 4 189 ASVs. According to our data, AM is dominated by the phyla Proteobacteria (mean relative abundance ± SD = 42.8 ± 19.4%) and Firmicutes (27.4 ± 18.9%), with Actinobacteria (14.8 ± 10.9%) and Bacteroidetes (9.2 ± 8.6%) being subdominant. At the family level, the most represented taxa are *Comamonadaceae* (6.1 ± 13.4%) and *Sphingomonadaceae* (4.3 ± 5.0%), both belonging to Proteobacteria. Other represented families are *Ruminococcaceae* (3.9 ± 7.6%), *Enterobacteriaceae* (3.7 ± 5.9%), *Clostridiaceae* (3.6 ± 6.8%), *Bacillaceae* (3.5 ± 5.0%) and *Flavobacteriaceae* (3.4 ± 5.7%). Please see **Supplementary Figure S2** for a graphical representation of the overall compositional structure of AM throughout the entire sampling period.

In order to explore connections between the AM structure and seasonality, we compared the levels of AM diversity over the different months (**Figure 3**). Diversity measurements indicated a general trend of microbial richness to decrease from winter to summer, although the differences did not reach statistical significance (Kruskal-Wallis test, FDR corrected p-value > 0.05) (**Figure 3A**). Conversely, the PCoA of unweighted UniFrac distances between the AM compositional profiles showed sample segregation according to the month of sampling (**Figure 3B**) (FDR-corrected permutation test with pseudo-F ratio, p-value = 0.012), meaning that seasonality significantly affects the overall compositional AM structure.

**Figure 3.**
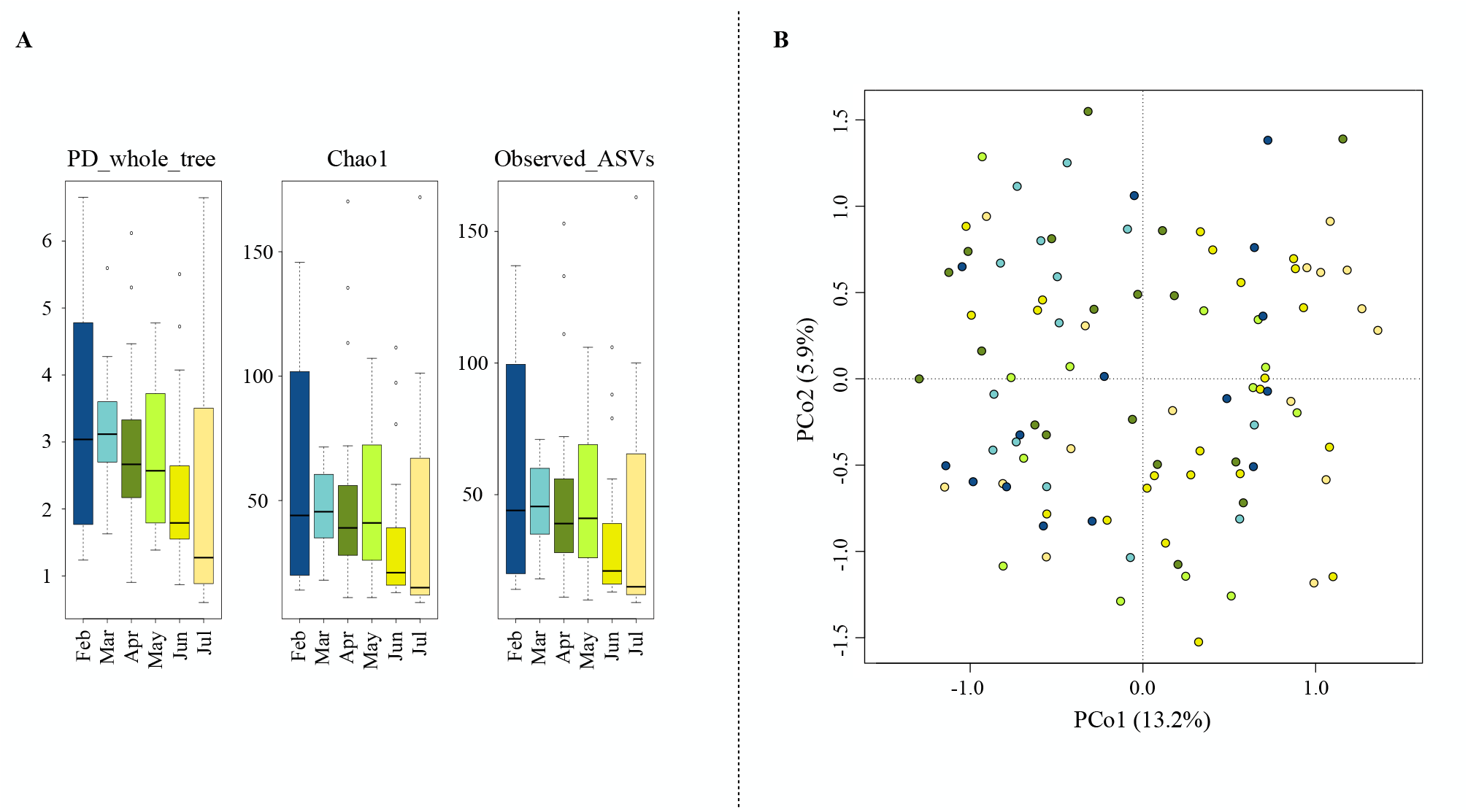
AM alpha and beta diversity throughout the sampling period. (**A**) Box-and-whiskers distribution of Faith’s Phylogenetic Diversity (PD_whole_tree), Chao1 index for microbial richness and number of observed ASVs, for each month of sampling. The data show a trend towards reduced microbial richness from winter to summer, although the differences did not reach statistical significance (Kruskal-Wallis test, FDR-corrected p-value > 0.05). (**B**) Principal Coordinates Analysis (PCoA) based on unweighted UniFrac distances between AM profiles, showing separation by sampling month (permutation test with pseudo-F ratio, p-value = 0.012) (same colour code as in panel A). The first and second principal components (PCo1 and PCo2) are plotted and the percentage of variance in the dataset explained by each axis is reported.

### Variation of the AM topological structure and association with PM emission sources and meteorological parameters

To further explore the overall AM variation across the sampling period, a clustering analysis of the AM compositional profiles was carried out. Hierarchical Ward-linkage clustering based on the Spearman correlation coefficients of family-level AM profiles resulted in the significant separation of 4 clusters, named C1, C2, C3 and C4, respectively (FDR-corrected permutation test with pseudo-F ratio, p-value ≤ 0.001) (**Figure 4**). Confirming the robustness of the identified clusters, the PCoA of the unweighted UniFrac distances between samples revealed a sharp segregation based on the assigned cluster (**Figure 5**). Interestingly, when we searched for correlations between PCoA coordinates and measured meteorological parameters or PMF factors (**Supplementary Tables S1 and S2**, respectively), we found that factor 5 (naval-maritime transport) and relative humidity (RH) were both positively correlated with the PCo1 axis (Kendall’s test, FDR-corrected p-value ≤ 0.001), while factor 6 (coal burning) was negatively correlated with the PCo1 coordinates (p-value ≤ 0.001) (**Figure 5**). This indirect gradient analysis allowed to highlight positive associations between clusters C1 and C3 and factors 6 and 5, respectively. Further, cluster C3 was found to be positively related to RH. As for seasonality, the clusters C3 and C4 are the most prevalent in summer and winter, respectively, while for C1 and C2 we did not observe any prevalence for a particular sampling period. We also compared the microbial diversity values of samples included in the different clusters, using three different diversity metrics. Our data indicated higher biodiversity in clusters C1 and C2 (PD whole tree, chao1, and observed ASVs, mean ± SD: 3.6±1.3, 67.2±43.9, and 65.0±40.5 for C1, 3.2±1.4, 59.1±38.2 and 57.6±36.0 for C2, respectively) compared to C3 and C4 (1.4±0.7, 18.7±9.4, and 18.5±9.3 for C3, 2.4±1.2, 31.2±18.4 and 31.2±18.3 for C4), with C3 having the lowest biodiversity (Kruskal-Wallis test, FDR corrected p-value ≤ 0.001).

**Figure 4.**
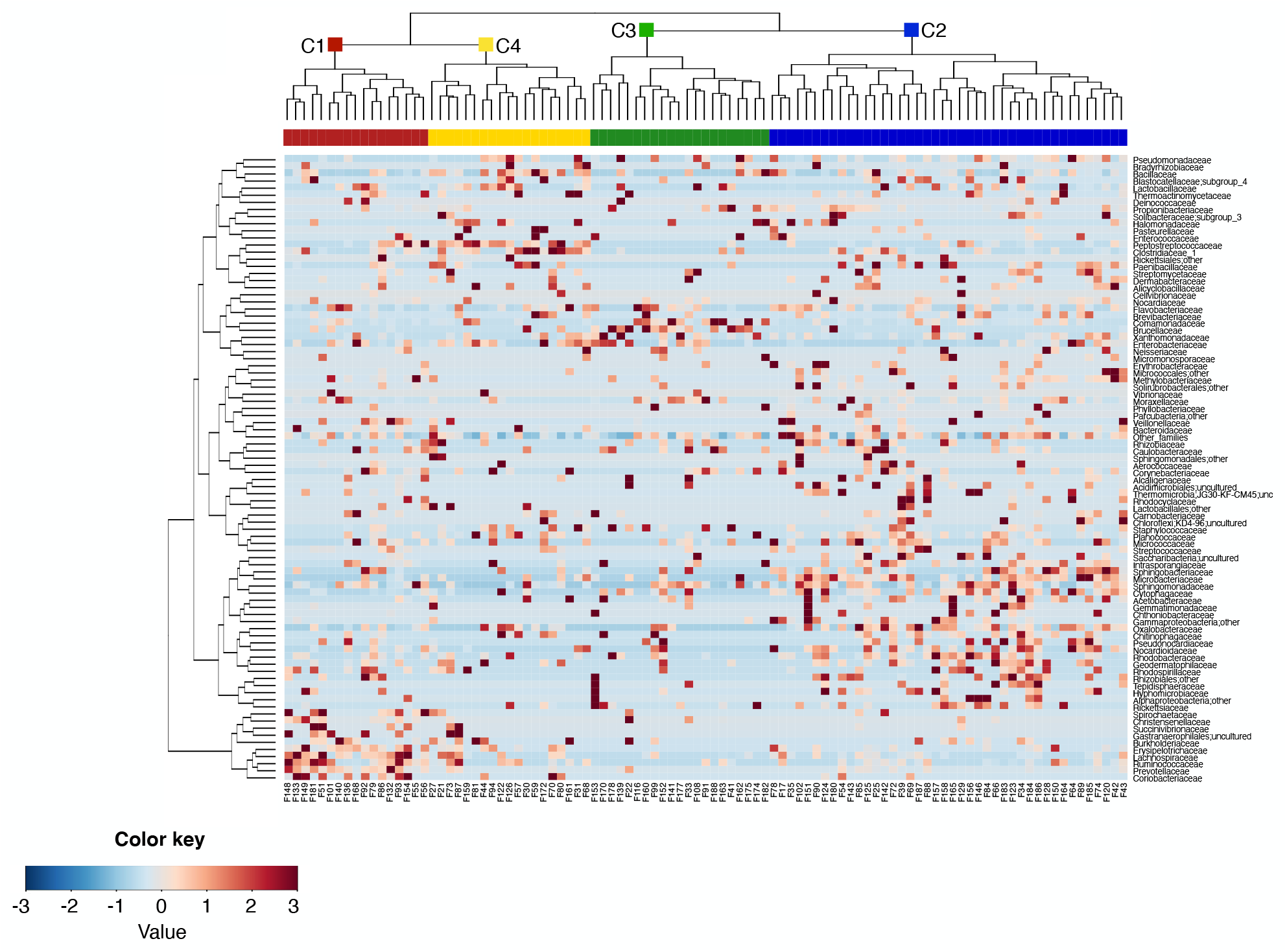
Family-level clusters of the airborne microbiome. Hierarchical Ward-linkage clustering based on the Spearman correlation coefficients of the proportion of families in the AM samples. Only families with relative abundance >2% in at least 3 samples were retained. The four identified clusters (FDR-corrected permutation test with pseudo-F ratio, p-value ≤ 0.001) are labelled in the top tree and highlighted by different coloured squares (red, blue, green and yellow for the clusters C1, C2, C3 and C4, respectively).

**Figure 5.**
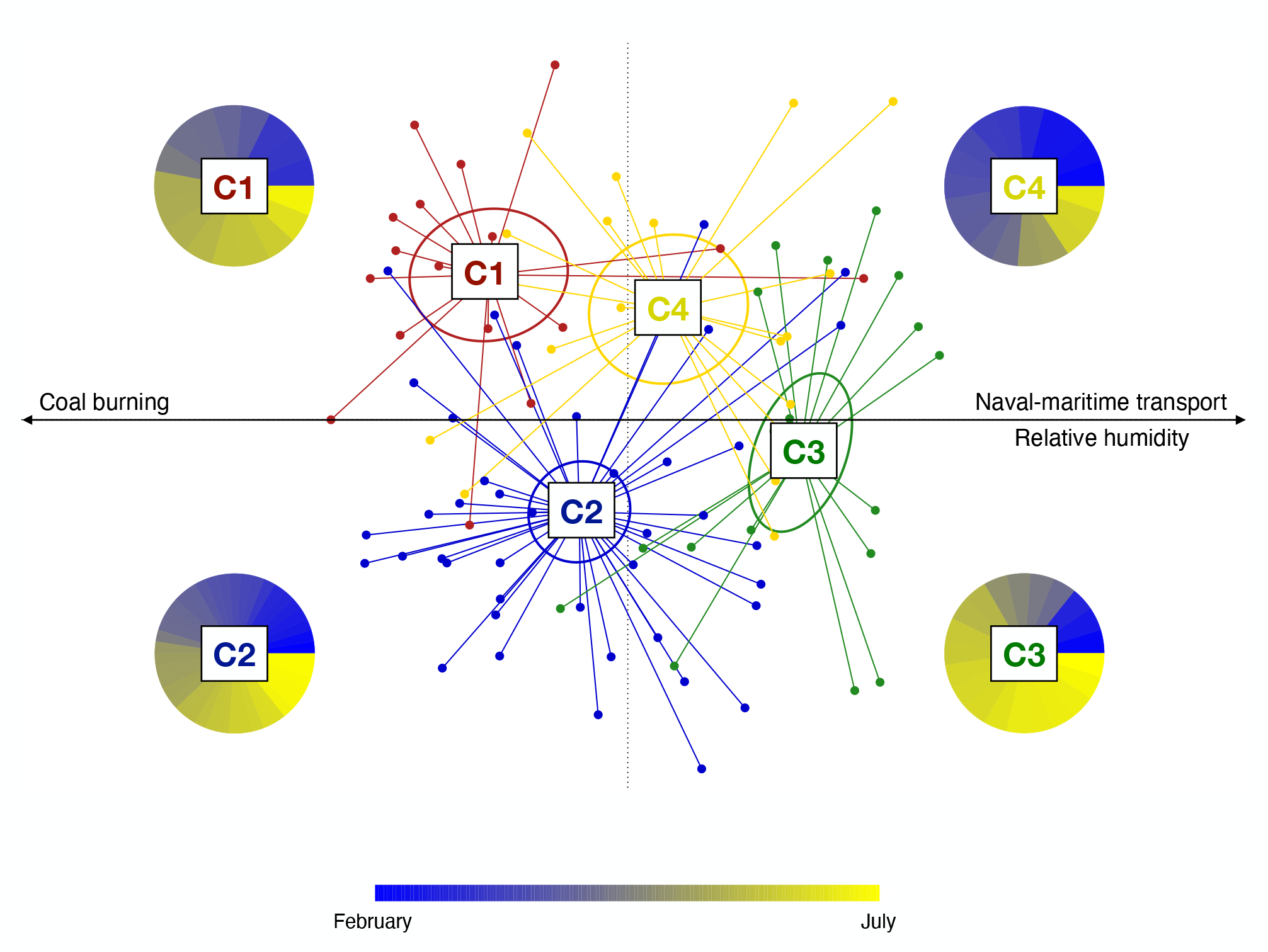
Variation of the AM topological structure and association with PM emission sources and meteorological parameters. Principal coordinates analysis (PCoA) based on the unweighted UniFrac distance shows separation between the microbial clusters (C1 to C4; permutation test with pseudo F-ratio, p-value ≤ 0.001; see also Figure 3). The percentage of variance in the dataset explained by each axis, first and second principal component (PCo1 and PCo2), is 13.2% and 5.9%, respectively. Ellipses include 95% confidence area based on the standard error of the weighted average of sample coordinates. Significant Kendall correlations between PCoA axes and PMF factors and measured meteorological parameters are reported with a black arrow. Specifically, the emission source factor 5 (naval-maritime transport) and relative humidity are both positively correlated with the PCo1 axis (Kendall correlation test, FDR-corrected p-value ≤ 0.001), while the emission source factor 6 (coal burning) is negatively correlated with the PCo1 coordinates (p-value ≤ 0.001). For each AM cluster, the proportion of samples based on the sampling time (from February (dark blue) to July (yellow) is shown as a pie chart.

### Compositional specificity and prevalent microbiological source of the four AM clusters

We subsequently compared the relative abundance of AM families among the four clusters in order to find out the most distinctive families of each of them (**Supplementary Figure S3**). According to our findings, the discriminating families (i.e. families with significantly different relative abundance, based on Kruskal-Wallis test) for the microbial cluster C1 are *Prevotellaceae*, *Erysipelotrichaceae*, *Coriobacteriaceae*, *Christensenellaceae*, *Lachnospiraceae*, *Ruminococcaceae*, and *Spirochaetaceae*. The microbial cluster C2 is instead characterized by higher abundance in the families *Microbacteriaceae*, *Cytophagaceae*, *Oxalobacteraceae*, *Sphingobacteriaceae*, *Nocardioidaceae*, *Methylobacteriaceae*, *Intrasporangiaceae*, *Rhodobacteraceae* and *Acetobacteraceae*. Only two proteobacterial families, namely *Brucellaceae* and *Comamonadaceae*, have a significantly higher abundance in cluster C3. Four families show higher abundance in cluster C4, i.e. *Peptostreptococcaceae*, *Clostridiaceae*, *Bacillaceae* and *Enterobacteriaceae*. It is also worth noting that the families *Planococcaceae* and *Paenibacillaceae* are highly represented in both C2 and C4 clusters, whereas *Sphingomonadaceae* members are equally represented in all clusters except for C4. In an attempt to identify the most likely prevalent microbial origin of the four AM clusters, we first derived the respective compositional peculiarities at the OTU level. To this aim, 16S rRNA gene reads were clustered at 97% homology, resulting in 3 821 OTUs. By linear regression, we subsequently obtained 80 OTUs specifically discriminating the four clusters. In particular, for 52 of these OTUs a significantly different distribution in the four clusters was confirmed by a Kruskal-Wallis test, as shown in **Supplementary Figure S4**. For each of them, the isolation source of the closest BLAST match within the NCBI 16S rRNA sequence database was recovered (**Supplementary Table S3**). Interestingly, according to our findings, the cluster C1 is mainly characterized by OTUs of faecal origin. These OTUs include sequences assigned to typical components of the human gut microbiome, such as *Faecalibacterium prausnitzii*, *Ruminococcus faecis*, *Prevotella copri*, *Eubacterium eligens*, *Ruminococcus bromii*, *Roseburia inulinivorans* and *Blautia faecis* [69–71], the cattle rumen components *Succinivibrio dextrinosolvens* [72] and *Oscillibacter ruminantium* [73], and the porcine gut microbiome member *Treponema porcinum* [74]. Differently, the cluster C2 is characterized by OTUs assigned to microorganisms isolated from plant roots and leaves, including *Curtobacterium flaccumfaciens* [75], *Glutamicibacter halophytocola* [76] and *Frigoribacterium endophyticum* [77], as well as by a specific pattern of environmental bacteria, from soil, air, and fresh and marine water ecosystems. Similarly, both clusters C3 and C4 are characterized by a peculiar combination of environmental microorganisms from different sources, including soil, fresh and marine waters, and airborne microbial ecosystems.

## Discussion

In order to explore connections between the local air microbiome, atmospheric pollution and meteorological factors, here we provide a longitudinal survey of the near-ground AM, atmospheric particulate and atmospheric parameters in Savona, Italy. According to our findings, the local AM appears dominated by the phyla Proteobacteria, Firmicutes, Actinobacteria and Bacteroidetes, well matching the general layout of an AM community [4]. The application of the PMF receptor modelling on the chemical compositional pattern of the PM_10_ samples collected during the field campaign allowed the identification of seven emission sources: “crustal material and road dust resuspension”, “traffic and biomass burning”, “secondary nitrate”, “secondary ammonium sulphate”, “naval-maritime transport”, “coal burning” and “sea spray”. Each source factor was subsequently subjected to anemological analysis based on polar plots, allowing each emission source to be associated with the corresponding wind direction to which the receptor site is downwind. Specifically, emission sources as “crustal material and road dust resuspension”, “secondary nitrate”, “secondary ammonium sulphate” and “coal burning” were associated with winds blowing from the inland toward the sampling site, intercepting traffic and industrial particulate sources. Conversely, emission sources such as “naval-maritime transport” and “sea spray” were associated with a sea breeze, supporting a marine origin for both. Finally, the “traffic and biomass burning” emission source mostly showed a local origin.

When we explored the AM structure variation during the observation period, we were able to identify four distinct clusters of samples, named C1 to C4. Interestingly, the four clusters were associated with a peculiar combination of seasonality, meteorological variables and emission sources. In particular, the AM cluster C1 was associated with the “coal burning” emission source, suggesting not actually the industrial facility as a microbiome source, but rather the influence of an air mass whose transport over a given district harvests chemical and microbiological components along the same tropospheric path. Instead, the cluster C3, most represented in the warm period, probably has a marine origin due to its association with the “naval-marine transport” emission source and high relative humidity. Finally, the clusters C2 and C4 did not show any specific association with the aerosol sources assessed by PMF, even if they showed a different seasonal behaviour, with C4 being more represented in the cold period.

The four AM clusters revealed a distinct, well-defined compositional structure, each being enriched with a specific set of microbial families and OTUs. The specificity of each bacterial profile de facto serves as a microbiological fingerprint, allowing to single out the probable microbiome sources characterizing each cluster that, similar to what occurs to abiotic particles, allow to trace back the origin of the air mass. In particular, the clusters C3 and C4 substantially reflect interconnected environmental microbiomes, encompassing a specific combination of microorganisms from soil resuspension, as well as from marine and fresh waters (possibly from rivers and streams flowing into the Ligurian Sea) and from the air. C2 cluster reveals the plant microbiome as an additional source, showing a further combination of plant-associated and environmental microorganisms, due to the contact of air masses over a vegetation landscape. Interestingly, the feasibility of air mass tracing also using bacterial species clearly emerges when we observe in detail the compositional structure of C1. This is in fact the only AM cluster carrying a recognizable pool of bacterial moieties of faecal origin, which are consistently part of the animal gut microbiome, suggesting not only a well-defined origin but also the potential use of this information in the assessment of microbiological impacts. It should be noted that in the area upwind C1 no sewage treatment plant as a possible source of faecal microbiome was present at the time of sampling. However, the area is very densely populated and forested areas populated by local fauna are closely found within a few kilometres.

Taken together, our data on the temporal dynamics of the near-ground AM in Savona, highlight the relevant degree of plasticity of AM over time. As such, we demonstrated how meteorological factors (e.g. wind direction and humidity) and atmospheric pollution (particles emission sources) can combine in shaping the AM configuration. In particular, coal burning and winds blowing from the inlands mix to establish a characteristic AM with a prevalence of aerosolized faecal microorganisms, regardless of seasonality. Conversely, in the summer season, humidity, sea breeze and naval-marine transport pollutants result in an AM mainly originating from environmental microbiomes, including microorganisms that are typically found in seawater and soil. Even if we were not able to establish connections between the other identified emission sources and specific AM clusters, we would stress the importance of seasonality in shaping the AM structure. Indeed, the variation between the clusters C2 and C4, for which no connection with any emission source was observed, was shown to be dependent on the sampling period, with the cluster C2 most prevalent during the warm season and including plant microbiomes as possible characteristic sources.

In conclusion, our results suggest that, in an urban settlement, air pollution may increase the proportion of aerosolized faecal microorganisms in the atmosphere, ultimately increasing citizens’ exposure to faecal microbes. Similar results have recently been obtained by exploring AM in Beijing over 6 months [22]. Our findings strengthen the importance of including the monitoring of the AM compositional structure as a determinant factor in the currently used air quality indexes. Indeed, in urban areas, the possible increased exposure to faecal-associated microbiomes as a result of increasing pollution can pose a possible threat to human health, particularly in regions with high-intensity animal farming, due to the inherent propensity of opportunistic pathogens to aerosolize.

## Acknowledgments

The authors wish to acknowledge Google Earth and the data providers SIO, NOAA, U.S. Navy, NGA, GEBCO and TerraMetrics for providing maps and 3D views of the sampling site.

This study represents partial fulfilment of the requirements for the PhD thesis of G. Palladino at the PhD course of Innovative Technologies and Sustainable Use of Mediterranean Sea Fishery and Biological Resources (FishMed – University of Bologna, Italy).

## Supplementary Information Description

**Supplementary Table S1 (2 pages) – Meteorological parameters during the PM sampling period.** The first column reports the sample ID, while the second indicates the sampling date. The meteorological parameters taken into account are temperature (T, °C), relative humidity (RH, %), pressure (P, mbar), rainfall (Rain, mm), wind speed (ws, m/s, and wind direction (wd, °). All values were taken every 30 min and averaged on a daily basis.

**Supplementary Table S2 (2 pages) – Normalized contributions per sample of the seven factors resolved by PMF analysis.** The first column reports the sample ID. All the other columns represent the contribution of each factor identified by PMF on the corresponding sample.

**Supplementary Table S3 (provided as Excel file) – Characteristics of the OTUs accounting for the compositional specificity of the four AM clusters.** For each OTU, the following information is given: unique OTUs ID, taxonomy as assigned with SILVA database, the cluster/s to which each OTU is significantly correlated (i.e. the cluster/s in which the given OTU is significantly more represented), the BLAST best hit resulting from blasting OTU fasta sequences against the NCBI 16S rRNA sequence database, the percentage of identity (ID (%)) and coverage (coverage (%)) between the OTU sequences and the corresponding best hit, and the isolation source of each best hit as reported in the GenBank database.

**Supplementary Figure S1 – Association between the factors obtained by PMF analysis and the wind direction and intensity.** Polar plots of the seven factors obtained by the PMF model. ws, wind speed; CPF, conditional probability function.

**Supplementary Figure S2 – AM overall composition.** Pie charts summarizing the microbiota composition of air filter samples at phylum (**A**) and family (**B**) level. Only phyla with relative abundance >1.5% in at least 10 samples and families with relative abundance >3% in at least 10 samples are shown.

**Supplementary Figure S3 – AM bacterial families differentially represented among the four microbial clusters.** Box plots showing the bacterial families whose relative abundance is significantly differently distributed among the microbial clusters C1-C4 (Kruskal-Wallis test, FDR-corrected p-value ≤ 0.05*, p-value ≤ 0.01** and p-value ≤ 0.001***). The central box represents the distance between the 25th and 75th percentiles. The median is marked with a black line. Whiskers identify the 10th and 90th percentiles.

**Supplementary Figure S4 (2 pages) – AM-associated OTUs showing different distribution across microbial clusters.** Box plots showing the OTUs whose relative abundance is significantly differently distributed among the four microbial clusters C1-C4 (Kruskal-Wallis test, FDR-corrected p-value ≤ 0.05*, p-value ≤ 0.01** and p-value ≤ 0.001***). The central box represents the distance between the 25th and 75th percentiles. The median is marked with a black line. Whiskers identify the 10th and 90th percentiles. unc., unclassified; amb., ambiguous.

