## Supplementary information for "Particulate Matter emission sources and meteorological parameters combine to shape the airborne microbiome communities in the Ligurian coast, Italy"

### SUPPLEMENTARY MATERIALS

**Supplementary Table S1 (2 pages) – Meteorological parameters during the PM sampling period.** The first column reports the sample ID, while the second indicates the sampling date. The meteorological parameters taken into account are temperature (T, °C), relative humidity (RH, %), pressure (P, mbar), rainfall (Rain, mm), wind speed (ws, m/s, and wind direction (wd, °). All values were taken every 30 min and averaged on a daily basis.

| Sample ID | Sampling date | T (°C) | RH (%) | P (mbar) | Rain (mm) | ws (m/s) | wd (°) |
|---|---|---|---|---|---|---|---|
| 1 | 01-Feb-2012 | 0.2 | 62.5 | 1019.3 | 0 | 4.6 | 318 |
| 2 | 05-Feb-2012 | -1.6 | 49.1 | 1029.3 | 0 | 2.3 | 316 |
| 3 | 06-Feb-2012 | -1.1 | 41.1 | 1023.5 | 0 | 3.7 | 321 |
| 4 | 09-Feb-2012 | 5.6 | 30.5 | 1025.5 | 0 | 1.9 | 323 |
| 5 | 11-Feb-2012 | -0.8 | 47.9 | 1024 | 0 | 3 | 323 |
| 6 | 14-Feb-2012 | 4.5 | 41.2 | 1020.3 | 0 | 0.9 | 315 |
| 7 | 15-Feb-2012 | 5.2 | 62.7 | 1016.6 | 0 | 0.5 | 242 |
| 8 | 17-Feb-2012 | 8.5 | 61.9 | 1027.2 | 0 | 0.5 | 219 |
| 9 | 18-Feb-2012 | 11.3 | 67.8 | 1026 | 0 | 0.8 | 237 |
| 10 | 19-Feb-2012 | 8.8 | 73.4 | 1023.7 | 0 | 0.1 | 270 |
| 11 | 23-Feb-2012 | 13.6 | 32.4 | 1028.4 | 0 | 1.9 | 318 |
| 12 | 25-Feb-2012 | 12.1 | 73.2 | 1026.6 | 0 | 0.2 | 197 |
| 13 | 26-Feb-2012 | 13 | 52.1 | 1019.3 | 0 | 0.6 | 35 |
| 14 | 27-Feb-2012 | 11.5 | 40.2 | 1026.3 | 0 | 0.7 | 318 |
| 15 | 28-Feb-2012 | 10.7 | 72.4 | 1028 | 0 | 0.4 | 232 |
| 16 | 06-Mar-2012 | 6.9 | 64.7 | 1023.5 | 1 | 3.7 | 314 |
| 17 | 09-Mar-2012 | 11.5 | 42.1 | 1035.4 | 0 | 4.2 | 306 |
| 18 | 10-Mar-2012 | 12.4 | 30.8 | 1034.6 | 0 | 3 | 324 |
| 19 | 11-Mar-2012 | 14.8 | 26.1 | 1026.9 | 0 | 2.5 | 317 |
| 20 | 12-Mar-2012 | 13.8 | 60 | 1025.3 | 0 | 0.5 | 334 |
| 21 | 14-Mar-2012 | 13.3 | 64.4 | 1031.5 | 0 | 0.8 | 326 |
| 22 | 19-Mar-2012 | 12.3 | 73 | 1028.7 | 1.4 | 0.7 | 150 |
| 23 | 21-Mar-2012 | 17 | 37.3 | 1037.4 | 0 | 3.4 | 313 |
| 24 | 23-Mar-2012 | 13.7 | 58.8 | 1031.8 | 0 | 0.4 | 92 |
| 25 | 24-Mar-2012 | 14.9 | 50.7 | 1028.5 | 0 | 0.5 | 358 |
| 26 | 25-Mar-2012 | 16 | 45 | 1029.2 | 0.8 | 0.7 | 350 |
| 27 | 27-Mar-2012 | 17.4 | 36.8 | 1031.3 | 0 | 0.7 | 260 |
| 28 | 28-Mar-2012 | 16.3 | 44.6 | 1029.7 | 0 | 0.3 | 79 |
| 29 | 29-Mar-2012 | 13.9 | 65.2 | 1024 | 0 | 0.4 | 76 |
| 30 | 02-Apr-2012 | 14.6 | 53.2 | 1017 | 0 | 1.9 | 330 |
| 31 | 03-Apr-2012 | 15 | 65.6 | 1015.2 | 0 | 0.5 | 67 |
| 32 | 04-Apr-2012 | 13.6 | 78.2 | 1014 | 23.4 | 0.3 | 307 |
| 33 | 05-Apr-2012 | 14.6 | 71.2 | 1013.7 | 4.6 | 0.8 | 279 |
| 34 | 08-Apr-2012 | 15.4 | 44.4 | 1006.9 | 0 | 0.9 | 42 |
| 35 | 09-Apr-2012 | 11.3 | 49.4 | 1017.1 | 0 | 0.3 | 303 |
| 36 | 10-Apr-2012 | 10.5 | 79.4 | 1015.9 | 5.2 | 0.1 | 61 |
| 37 | 11-Apr-2012 | 10 | 68.9 | 1006.6 | 18.6 | 1.6 | 313 |
| 38 | 12-Apr-2012 | 11.6 | 75.4 | 1009.5 | 0 | 0.3 | 115 |
| 39 | 13-Apr-2012 | 14.8 | 59.9 | 1005.5 | 8.8 | 1 | 304 |
| 40 | 14-Apr-2012 | 12.5 | 68.3 | 1001.5 | 0.6 | 1.5 | 306 |
| 41 | 15-Apr-2012 | 12.7 | 70.2 | 1003.3 | 0.8 | 0.4 | 38 |
| 42 | 16-Apr-2012 | 14 | 57.6 | 1009.7 | 1.2 | 1.2 | 305 |
| 43 | 17-Apr-2012 | 15.5 | 50.2 | 1011.7 | 0.2 | 0.6 | 272 |
| 44 | 18-Apr-2012 | 13.6 | 70.3 | 1004.1 | 0.6 | 0.5 | 95 |
| 45 | 23-Apr-2012 | NA | NA | 1012.7 | 0 | 0.1 | 270 |
| 46 | 25-Apr-2012 | 11.5 | 80.7 | 1016.4 | 0 | 0.3 | 252 |

| Sample ID | Sampling date | T (°C) | RH (%) | P (mbar) | Rain (mm) | ws (m/s) | wd (°) |
|---|---|---|---|---|---|---|---|
| 47 | 26-Apr-2012 | NA | NA | 1024.2 | 0 | NA | NA |
| 48 | 02-May-2012 | 14.2 | 79.7 | 1022.8 | 0.2 | 0.3 | 68 |
| 49 | 10-May-2012 | 18.1 | 71.9 | 1029.4 | 0 | 0 | 68 |
| 50 | 14-May-2012 | 16.3 | 51.3 | 1020.1 | 0 | 1.2 | 287 |
| 51 | 16-May-2012 | 18.2 | 29.6 | 1017.4 | 0 | 0.7 | 56 |
| 52 | 17-May-2012 | 16.9 | 27.3 | 1022.9 | 0 | 0.5 | 268 |
| 53 | 18-May-2012 | 16.2 | 51.4 | 1021.4 | 0 | 0.1 | 354 |
| 54 | 19-May-2012 | 15.8 | 54.2 | 1020.8 | 0.4 | 0.1 | 277 |
| 55 | 20-May-2012 | 14.8 | 71.3 | 1017.5 | 11.4 | 0.3 | 291 |
| 56 | 22-May-2012 | 17.4 | 59.6 | 1012.3 | 0 | 0.2 | 256 |
| 57 | 23-May-2012 | 18.2 | 72.9 | 1019 | 0 | 0.1 | 68 |
| 58 | 26-May-2012 | 20.8 | 56.6 | 1021.4 | 0 | 0.2 | 276 |
| 59 | 27-May-2012 | 20.5 | 64 | 1022.1 | 0 | 0.1 | 149 |
| 60 | 30-May-2012 | 20.4 | 75.5 | 1022.9 | 0 | 0.3 | 236 |
| 61 | 02-Jun-2012 | 19.7 | 85.8 | 1021.2 | 1.2 | 0.3 | 79 |
| 62 | 03-Jun-2012 | 20 | 86.8 | 1019.9 | 5.4 | 0.5 | 73 |
| 63 | 04-Jun-2012 | 20.4 | 77 | 1013.4 | 1 | 0.4 | 181 |
| 64 | 05-Jun-2012 | 19.5 | 71.1 | 1016.9 | 0 | 0.3 | 68 |
| 65 | 06-Jun-2012 | 18.6 | 78.6 | 1018.6 | 0 | 0.2 | 68 |
| 66 | 09-Jun-2012 | 21.3 | 75.3 | 1017.3 | 0 | 0.1 | 231 |
| 67 | 11-Jun-2012 | 20.8 | 74.7 | 1009.6 | 0 | 0.2 | 69 |
| 68 | 12-Jun-2012 | 19.6 | 72.8 | 1007.7 | 0 | 0.6 | 64 |
| 69 | 13-Jun-2012 | 19.7 | 71.1 | 1016.6 | 0 | 0.6 | 67 |
| 70 | 14-Jun-2012 | 20 | 69.9 | 1023.3 | 0 | 0.7 | 69 |
| 71 | 15-Jun-2012 | 20 | 62.7 | 1026.1 | 0 | 0.2 | 68 |
| 72 | 16-Jun-2012 | 25.3 | 51.7 | 1025.1 | 0 | 0.2 | 36 |
| 73 | 17-Jun-2012 | 24.2 | 55.3 | 1023.5 | 0 | 0.1 | 68 |
| 74 | 19-Jun-2012 | 25.8 | 61.8 | 1020.7 | 0 | 0.1 | 63 |
| 75 | 20-Jun-2012 | 24.6 | 58.3 | 1018.5 | 0 | 0.1 | 293 |
| 76 | 21-Jun-2012 | 23.3 | 66.9 | 1015.9 | 0 | 0.4 | 63 |
| 77 | 22-Jun-2012 | 24.1 | 71.4 | 1019.9 | 0 | 0.3 | 66 |
| 78 | 23-Jun-2012 | 24.9 | 69.3 | 1022.9 | 0 | 0.2 | 85 |
| 79 | 24-Jun-2012 | 24.8 | 70.2 | 1022.4 | 0 | 0.2 | 80 |
| 80 | 25-Jun-2012 | 23.6 | 74.9 | 1018 | 0 | 0.2 | 68 |
| 81 | 26-Jun-2012 | 25.2 | 66.2 | 1019.8 | 0 | 0.1 | 62 |
| 82 | 27-Jun-2012 | 28.6 | 49.1 | 1021.3 | 0 | 0.3 | 14 |
| 83 | 28-Jun-2012 | 26.6 | 58.5 | 1019.3 | 0 | 0.2 | 68 |
| 84 | 01-Jul-2012 | 25.8 | 71.2 | 1019.2 | 0 | 0.2 | 59 |
| 85 | 03-Jul-2012 | 23.4 | 68.8 | 1020.3 | 0 | 0.3 | 68 |
| 86 | 05-Jul-2012 | 24.5 | 61 | 1016.3 | 0 | 0.1 | 60 |
| 87 | 07-Jul-2012 | 23.9 | 76.4 | 1018.4 | 0.6 | 0.6 | 66 |
| 88 | 08-Jul-2012 | 24.4 | 73.5 | 1017.8 | 0 | 0.3 | 69 |
| 89 | 10-Jul-2012 | 25.4 | 71.9 | 1017.6 | 0 | 0.4 | 72 |
| 90 | 11-Jul-2012 | 24.9 | 74.3 | 1018.4 | 0.2 | 0.4 | 68 |
| 91 | 13-Jul-2012 | 23.6 | 69.7 | 1015 | 0 | 0.3 | 68 |
| 92 | 14-Jul-2012 | 23.7 | 76.5 | 1013.6 | 0 | 0.8 | 71 |
| 93 | 15-Jul-2012 | 23.5 | 68.4 | 1015.7 | 0 | 0.3 | 68 |
| 94 | 16-Jul-2012 | 25.7 | 39.8 | 1024 | NA | 0.8 | 281 |
| 95 | 17-Jul-2012 | 25.4 | 44 | 1025.9 | NA | 0.3 | 78 |
| 96 | 18-Jul-2012 | 24.8 | 55 | 1023.6 | NA | 0.3 | 69 |
| 97 | 19-Jul-2012 | 21.3 | 76.7 | 1021 | NA | 0 | 68 |
| 98 | 20-Jul-2012 | 0.2 | 62.5 | 1019.3 | 0 | 4.6 | 318 |

**Supplementary Table S2 (2 pages) –Normalized contributions per sample of the seven factors resolved by PMF analysis.** The first column reports the sample ID. All the other columns represent the contribution of each factor identified by PMF on the corresponding sample.

| Sample ID | Factor 1 | Factor 2 | Factor 3 | Factor 4 | Factor 5 | Factor 6 | Factor 7 |
|---|---|---|---|---|---|---|---|
| 1 | -0.01 | -0.12 | 2.37 | 1.7 | -0.2 | 2.96 | -0.2 |
| 2 | 0.03 | -0.2 | 3.73 | 1.17 | 0.18 | 3.04 | -0.14 |
| 3 | 2.83 | 0.47 | 3.42 | 1.22 | 0.13 | 4.42 | -0.2 |
| 4 | 4.17 | 1.34 | 2.84 | 1.24 | 0.31 | 5 | 0.18 |
| 5 | 0.89 | -0.2 | 4.4 | 3.68 | -0.2 | 3.34 | 0 |
| 6 | 0.94 | 0.96 | 0.98 | 1.96 | -0.16 | 1.2 | 0.24 |
| 7 | 0.16 | 1.74 | 0.9 | 0.9 | 0.29 | 0.54 | 0.71 |
| 8 | 1.25 | 0.97 | 3.39 | -0.2 | 1.21 | 0.55 | 0.59 |
| 9 | -0.2 | 0.93 | 2.79 | 0.18 | 1.6 | 0.8 | 0.88 |
| 10 | 0.67 | -0.2 | 0.28 | -0.09 | 1.01 | 9.02 | -0.07 |
| 11 | 2.29 | 0.09 | -0.02 | -0.2 | 0.79 | 5.42 | -0.08 |
| 12 | 0.31 | 0.36 | 2.42 | 0.48 | 2.25 | 1.42 | 0.1 |
| 13 | 0.61 | 1.23 | 0.41 | 0.93 | 0.98 | 1.61 | -0.01 |
| 14 | 1.9 | 2.14 | 0.3 | 0.11 | 0.4 | 2.38 | -0.14 |
| 15 | 0.02 | 2.57 | 2.02 | 0.68 | 0.67 | 0 | 0.2 |
| 16 | -0.05 | 0.36 | 1 | 0.65 | -0.04 | 1.29 | -0.09 |
| 17 | 1.74 | 0.28 | 3.35 | 0.39 | 0.14 | 0.9 | -0.09 |
| 18 | 1.89 | 0.17 | 2.61 | 0.52 | 0.44 | 1.38 | -0.13 |
| 19 | 0.75 | 0.38 | 3.12 | 0.34 | 0.57 | 1.39 | -0.08 |
| 20 | 1.03 | 1.98 | 3.08 | 0.17 | 0.97 | 1.21 | 0.03 |
| 21 | 0.08 | 1.7 | 8.18 | 0.63 | 0.18 | 0.4 | 0.06 |
| 22 | 0.08 | 1.38 | 0.37 | 0.17 | 0.85 | 0.33 | 3.16 |
| 23 | 2.76 | 1.06 | 0.18 | 0.58 | 0.49 | 1.94 | -0.2 |
| 24 | 1.58 | 3.06 | 1.85 | 1.71 | 0.54 | 2.5 | -0.1 |
| 25 | 2.27 | 2.17 | 0.76 | 2.56 | 0.29 | 0.49 | 0.04 |
| 26 | 3.26 | 0.93 | 1.15 | 1.32 | 0.58 | 0.96 | 0.15 |
| 27 | 2.36 | 1.71 | -0.2 | 1.7 | -0.05 | 1.98 | -0.17 |
| 28 | 1.71 | 3.84 | -0.16 | 1.53 | -0.2 | 3.08 | -0.14 |
| 29 | 1.24 | 1.52 | 1.83 | 2.36 | 0.34 | 1.53 | -0.03 |
| 30 | 4.01 | 0.15 | 1.05 | -0.2 | 2.21 | 3.42 | 0.64 |
| 31 | 2.65 | 0.27 | 1.74 | 0.04 | 2.18 | 1.61 | 0.31 |
| 32 | 0.68 | -0.1 | 4.53 | 1.13 | 0.46 | 1.42 | 0 |
| 33 | 0.12 | 0.99 | 0.41 | 0.51 | 0.26 | 0.11 | 0.13 |
| 34 | 1.32 | 0.35 | 0.08 | 0.39 | 0.12 | 0.3 | 0.43 |
| 35 | 0.54 | 0.72 | 0.53 | 0.34 | 0.41 | 0.29 | 0.65 |
| 36 | -0.18 | 1.84 | 1.21 | 0.72 | 0.38 | 0.9 | 0.07 |
| 37 | 0.18 | 0.84 | -0.16 | 0.09 | -0.04 | 0.94 | -0.05 |
| 38 | 0.17 | 0.79 | 0.7 | -0.18 | 0.88 | 0.42 | 2.75 |
| 39 | 0.49 | 0.42 | 0.39 | 0.32 | 0.63 | 0.74 | 0.24 |
| 40 | -0.13 | 0.57 | 0.01 | 0.4 | -0.04 | 0.33 | -0.01 |
| 41 | -0.19 | 0.59 | 0.21 | 0.37 | 0.17 | 0.25 | 0.22 |
| 42 | 0.54 | 0.68 | 0.04 | 0.46 | -0.05 | 0.3 | -0.01 |
| 43 | 0.24 | 1.08 | 0.28 | 0.51 | -0.1 | 0.75 | 0.33 |
| 44 | 0.18 | 1.94 | 0.83 | 0.01 | 0.81 | 1.26 | 3.35 |
| 45 | 0.14 | 0.66 | 0.09 | 0 | -0.07 | 0.37 | 8.38 |
| 46 | 0.21 | 0.38 | 0.27 | -0.12 | 0.79 | 0.13 | 6.81 |
| 47 | 0.26 | 0.6 | 0.61 | -0.06 | 1.3 | 0.94 | 3.99 |

| Sample ID | Factor 1 | Factor 2 | Factor 3 | Factor 4 | Factor 5 | Factor 6 | Factor 7 |
|---|---|---|---|---|---|---|---|
| 48 | 0.57 | 0.52 | 0.78 | 0.32 | 1.47 | 1.01 | 0.35 |
| 49 | 0.09 | 1.91 | 0.2 | 1.1 | 2.04 | 1.1 | 0.03 |
| 50 | 0.74 | 1.13 | 0.77 | 0.73 | 0.81 | -0.2 | 0.82 |
| 51 | 3.12 | 1.29 | -0.2 | 0.33 | 0.13 | 0.38 | 0 |
| 52 | 1.45 | 0.79 | -0.1 | 0.4 | 0.26 | 0.73 | 0.08 |
| 53 | 1.62 | 2.78 | 0.21 | 0.34 | 0.31 | 0.59 | -0.03 |
| 54 | 2.19 | 1.91 | 1.53 | 0.76 | 0.02 | 0.14 | -0.02 |
| 55 | 0.74 | 0.5 | 4.04 | 0.58 | 0.11 | -0.2 | 0.07 |
| 56 | 0.29 | 1.16 | 0.13 | 0.13 | 0.17 | 0.74 | 0.78 |
| 57 | 0.02 | 1.74 | 1.17 | 0.05 | 1.48 | 0.29 | 0.41 |
| 58 | 0.95 | 1.6 | 0.19 | 1.76 | 0.04 | 1.43 | -0.09 |
| 59 | 0.25 | 1.47 | 0.18 | 1.9 | -0.14 | 1.32 | -0.08 |
| 60 | -0.2 | 1.22 | -0.06 | 2.94 | 0.44 | 0.7 | -0.03 |
| 61 | 0.31 | 0.31 | -0.2 | 1.71 | 0.62 | 1.08 | -0.04 |
| 62 | -0.16 | -0.2 | -0.2 | 3.74 | 2.36 | 0.79 | 0.13 |
| 63 | 0.13 | 0.85 | 0.23 | 0.94 | 1.11 | 0.07 | 2.08 |
| 64 | 0.11 | 0.83 | 1.31 | 0.08 | 1.99 | 0.05 | 1.68 |
| 65 | 0.34 | 0.53 | 0.57 | -0.19 | 2.85 | 0.05 | 0.73 |
| 66 | 0.07 | 0.22 | 1.65 | 0.53 | 2.55 | 0.28 | 4.15 |
| 67 | 0.29 | 0.5 | 0.37 | 0.47 | 2.27 | 0.36 | 6.23 |
| 68 | 0.39 | 0.48 | -0.2 | 0.2 | 0.62 | 0.27 | 8.04 |
| 69 | 0.17 | 0.31 | 0.53 | -0.2 | 2.2 | 0.15 | 7.55 |
| 70 | 0.05 | -0.2 | 1.14 | 0.18 | 1.92 | 0.18 | 5.54 |
| 71 | 0.31 | 1.37 | 0.66 | 0.44 | 2.36 | 0.48 | 0.11 |
| 72 | 1.23 | 1.38 | 0.3 | 1.66 | 1.22 | 0.65 | -0.01 |
| 73 | 0.45 | 2.44 | 0.27 | 1.88 | 0.72 | 0.18 | 0.04 |
| 74 | 1.87 | 2.97 | 0.22 | 2.89 | 0.06 | 0.24 | 0 |
| 75 | 2.48 | 2.24 | -0.11 | 2.57 | 0.1 | 1.39 | -0.04 |
| 76 | 3.19 | 0.44 | 0.84 | 1.61 | 3.58 | 0.29 | 0.28 |
| 77 | 3.34 | 0.38 | -0.1 | 3.68 | 2.98 | 0.62 | -0.06 |
| 78 | 1.3 | 0.68 | 0.44 | 3.58 | 3.21 | 0.75 | 0.05 |
| 79 | 1.12 | 1.1 | -0.02 | 3.3 | 2.47 | 0.92 | 0.01 |
| 80 | 0.13 | 0.74 | 0.88 | 2.93 | 2.8 | -0.01 | 0.06 |
| 81 | 0.85 | 2.22 | 0.23 | 2.4 | 2.13 | 0.11 | 0.02 |
| 82 | 2.44 | 1.31 | -0.16 | 0.23 | 1.18 | 1.02 | -0.13 |
| 83 | 0.73 | 1.96 | 0.37 | 2.24 | 0.72 | 0.37 | -0.02 |
| 84 | 3.45 | 0.53 | -0.18 | 2.88 | 1.04 | 0.31 | -0.11 |
| 85 | 1.6 | 0 | 1.45 | 0.92 | 1.27 | 0.53 | 3.41 |
| 86 | 2.4 | 1.89 | 0.08 | 1.78 | 0.36 | 0.22 | -0.06 |
| 87 | 0.77 | 0.22 | 0.75 | -0.16 | 3.71 | 0.55 | 0.34 |
| 88 | 0.48 | 0.76 | 0.01 | 1.72 | 1.81 | 0.2 | 0.01 |
| 89 | 0.19 | -0.01 | 1.21 | 1.13 | 2.83 | -0.2 | 0.38 |
| 90 | 0.33 | 0.06 | 1.22 | 1.24 | 3.15 | 0.26 | 0.45 |
| 91 | 0.28 | 0.53 | 1.51 | -0.17 | 1.29 | -0.09 | 9.64 |
| 92 | 0.24 | -0.01 | 1.75 | 1.49 | 0.83 | -0.02 | 5.21 |
| 93 | 0.43 | 0.45 | 0.74 | -0.05 | 2.12 | 0.07 | 6.8 |
| 94 | 3.51 | 1.76 | -0.03 | 0.42 | 0.44 | 0.63 | 0.36 |
| 95 | 2.07 | 2.15 | 0.02 | 0.62 | 0.51 | 0.6 | 0 |
| 96 | 1.37 | 2.47 | 0.11 | 1.42 | 0.36 | 0.48 | -0.05 |
| 97 | 0.03 | 0.76 | 0.98 | 1.54 | 2.1 | -0.2 | 0.19 |
| 98 | 0.2 | 0.12 | 0.45 | 1.93 | 2.33 | 0.76 | 0.07 |

**Supplementary Table S3 (provided as Excel file) – Characteristics of the OTUs accounting for the compositional specificity of the four AM clusters**. For each OTU, the following information is given: unique OTUs ID, taxonomy as assigned with SILVA database, the cluster/s to which each OTU is significantly correlated (i.e. the cluster/s in which the given OTU is significantly more represented), the BLAST best hit resulting from blasting OTU fasta sequences against the NCBI 16S rRNA sequence database, the percentage of identity (ID (%)) and coverage (coverage (%)) between the OTU sequences and the corresponding best hit, and the isolation source of each best hit as reported in the GenBank database.

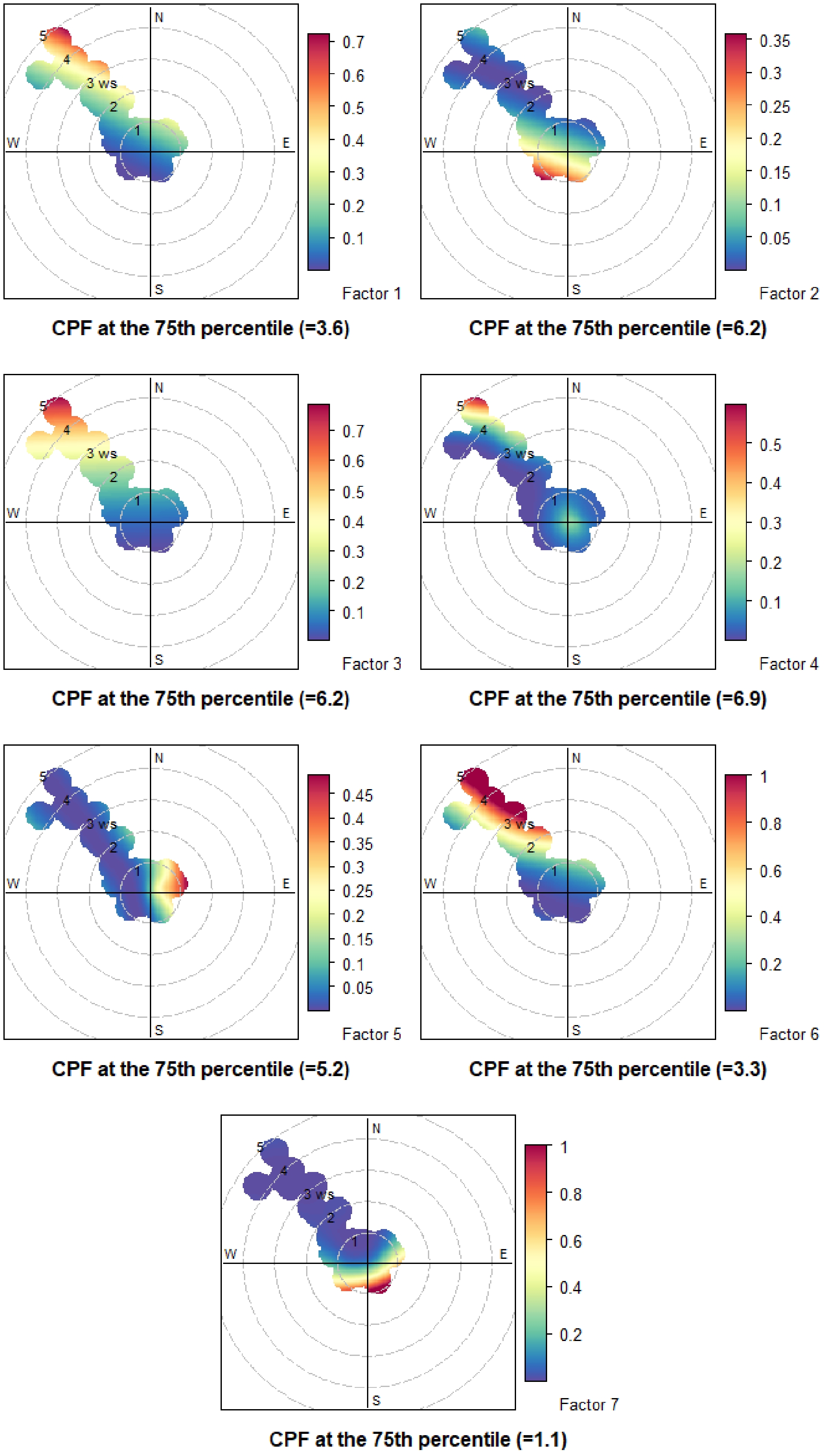


**Supplementary Figure S1 – Association between the factors obtained by PMF analysis and the wind direction and intensity.** Polar plots of the seven factors obtained by the PMF model. ws, wind speed; CPF, conditional probability function.

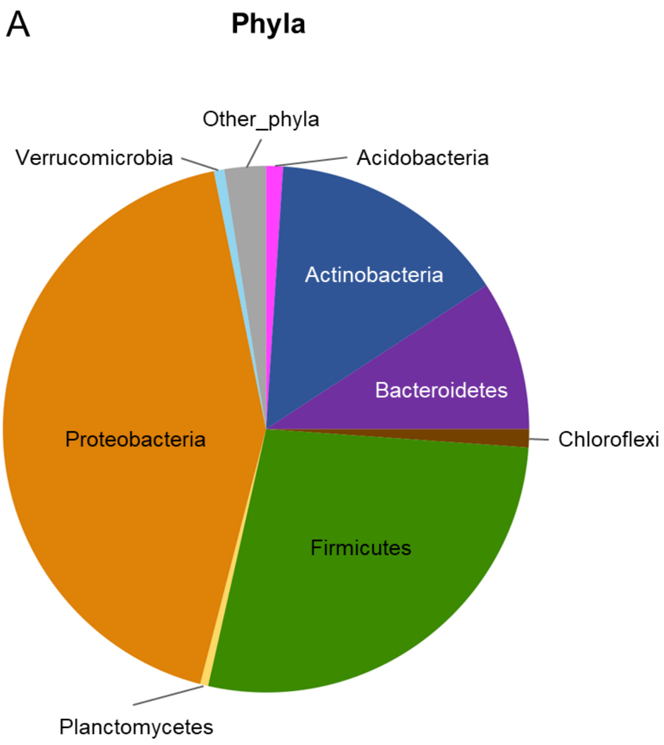


**Supplementary Figure S2 - AM overall composition.** Pie charts summarizing the microbiota composition of air filter samples at phylum (A) and family (B) level. Only phyla with relative abundance >1.5% in at least 10 samples and families with relative abundance >3% in at least 10 samples are shown.

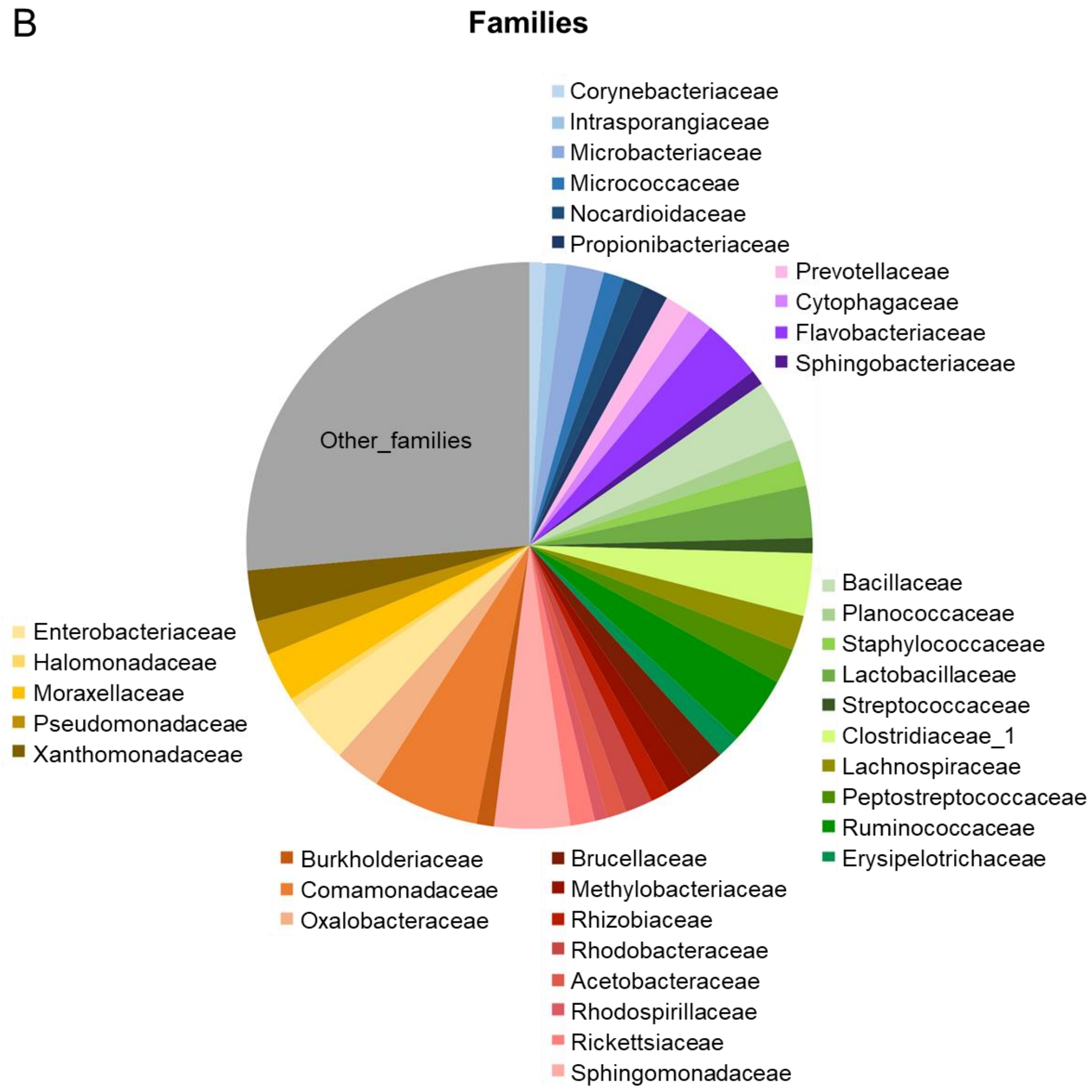

**Supplementary Figure S3 – AM bacterial families differentially represented among the four microbial clusters.** Box plots showing the bacterial families whose relative abundance is significantly differently distributed among the microbial clusters C1-C4 (Kruskal-Wallis test, FDR-corrected p-value ≤ 0.05*, p-value ≤ 0.01** and p-value ≤ 0.001***). The central box represents the distance between the 25th and 75th percentiles. The median is marked with a black line. Whiskers identify the 10th and 90th percentiles.

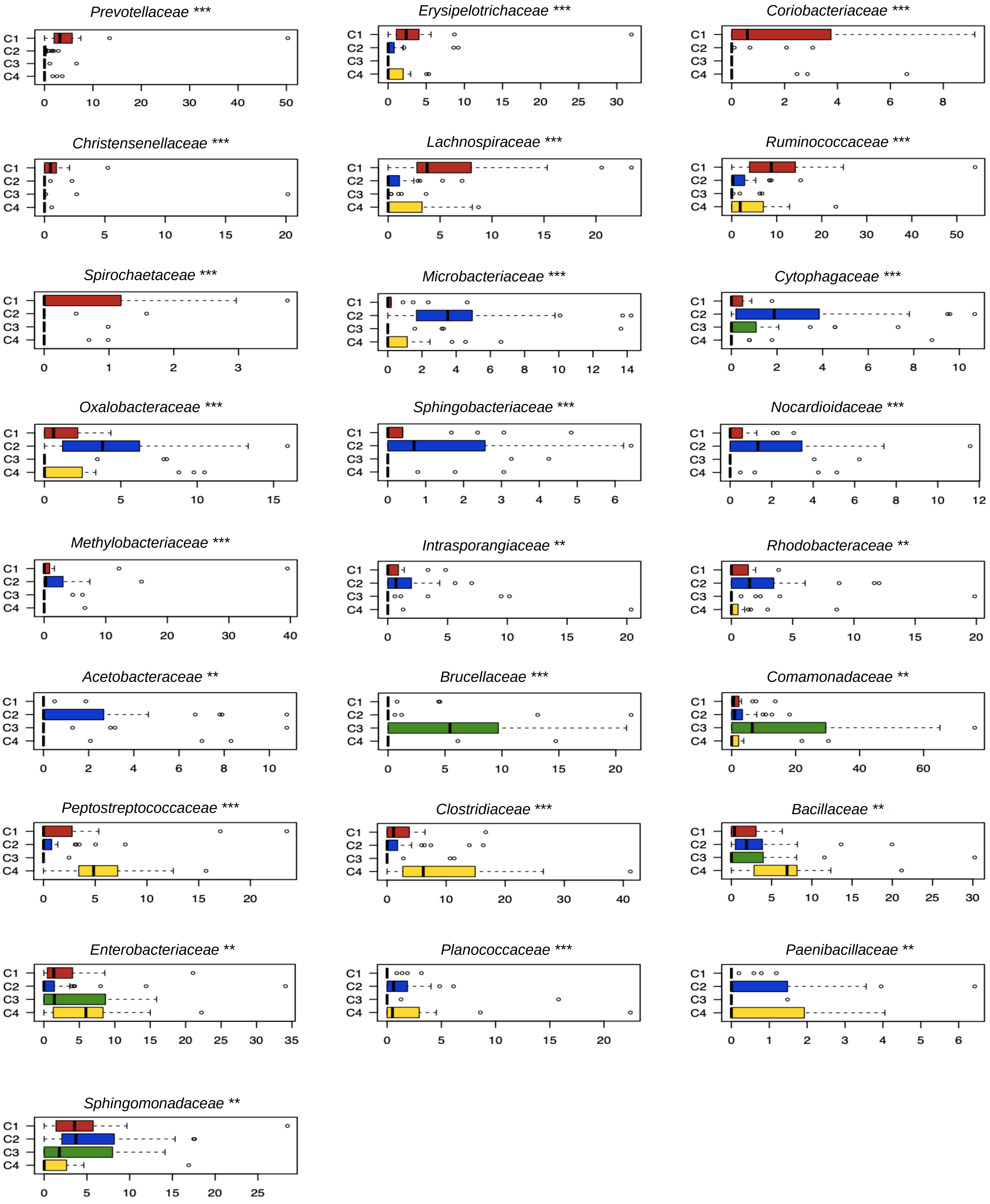

**Supplementary Figure S4 (2 pages) – AM-associated OTUs showing different distribution across microbial clusters.** Box plots showing the OTUs whose relative abundance is significantly differently distributed among the four microbial clusters C1-C4 (Kruskal-Wallis test, FDR-corrected p-value ≤ 0.05*, p-value ≤ 0.01** and p-value ≤ 0.001***). The central box represents the distance between the 25th and 75th percentiles. The median is marked with a black line. Whiskers identify the 10th and 90th percentiles. unc., unclassified; amb., ambiguous.

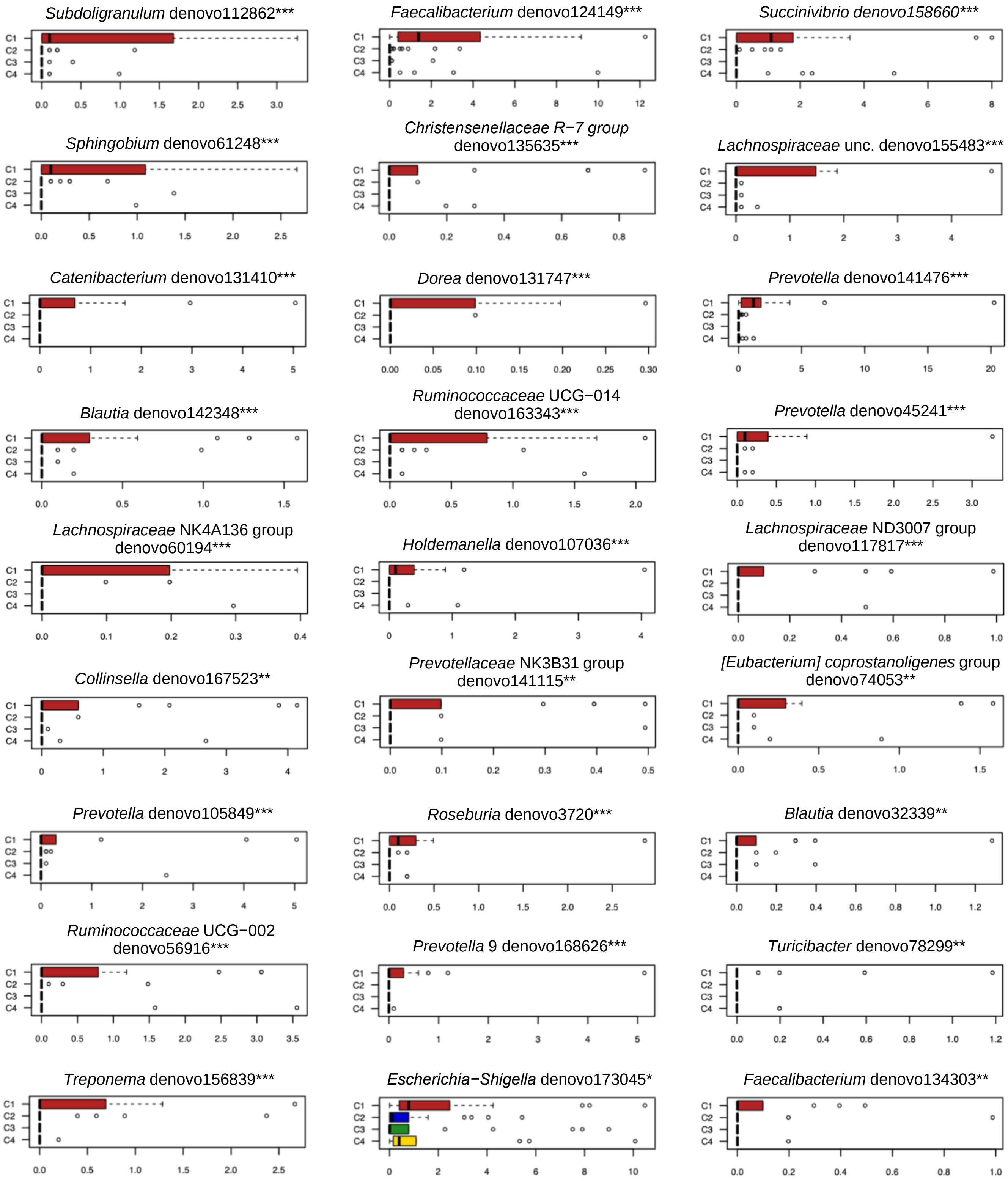

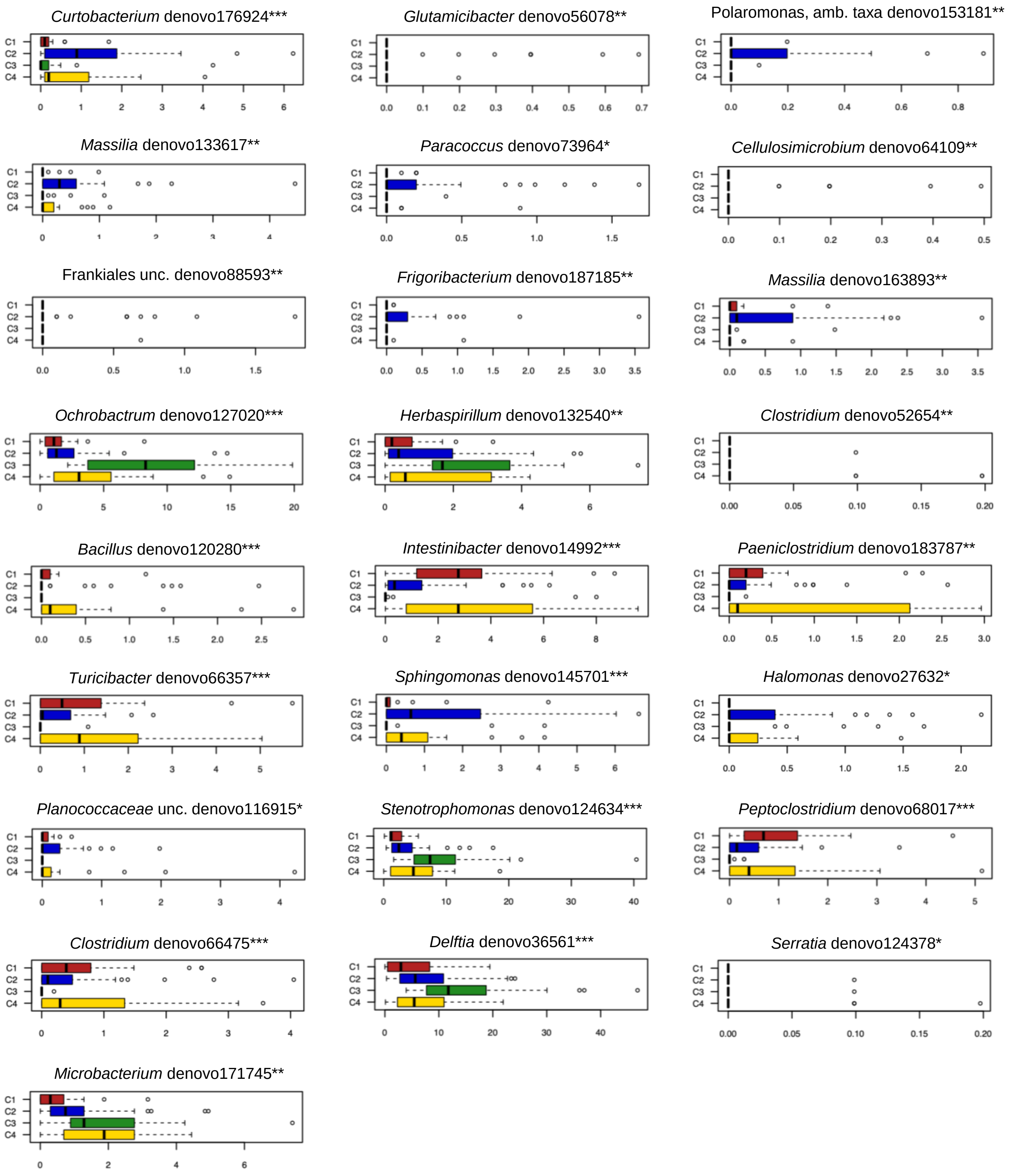
*Curtobacterium* denovo176924***
C1 C2 C3 C4
0 1 2 3 4 5 6
*Glutamicibacter* denovo56078**
C1 C2 C3 C4
0.0 0.1 0.2 0.3 0.4 0.5 0.6 0.7
Polaromonas, amb. taxa denovo153181**
C1 C2 C3 C4
0.0 0.2 0.4 0.6 0.8
*Massilia* denovo133617**
C1 C2 C3 C4
0 1 2 3 4
*Paracoccus* denovo73964*
C1 C2 C3 C4
0.0 0.5 1.0 1.5
*Cellulosimicrobium* denovo64109**
C1 C2 C3 C4
0.0 0.1 0.2 0.3 0.4 0.5
Frankiales unc. denovo88593**
C1 C2 C3 C4
0.0 0.5 1.0 1.5
*Frigoribacterium* denovo187185**
C1 C2 C3 C4
0.0 0.5 1.0 1.5 2.0 2.5 3.0 3.5
*Massilia* denovo163893**
C1 C2 C3 C4
0.0 0.5 1.0 1.5 2.0 2.5 3.0 3.5
*Ochrobactrum* denovo127020***
C1 C2 C3 C4
0 5 10 15 20
*Herbaspirillum* denovo132540**
C1 C2 C3 C4
0 2 4 6
*Clostridium* denovo52654**
C1 C2 C3 C4
0.00 0.05 0.10 0.15 0.20
*Bacillus* denovo120280***
C1 C2 C3 C4
0.0 0.5 1.0 1.5 2.0 2.5
*Intestinibacter* denovo14992***
C1 C2 C3 C4
0 2 4 6 8
*Paeniclostridium* denovo183787**
C1 C2 C3 C4
0.0 0.5 1.0 1.5 2.0 2.5 3.0
*Turicibacter* denovo66357***
C1 C2 C3 C4
0 1 2 3 4 5
*Sphingomonas* denovo145701***
C1 C2 C3 C4
0 1 2 3 4 5 6
*Halomonas* denovo27632*
C1 C2 C3 C4
0.0 0.5 1.0 1.5 2.0
*Planococcaceae* unc. denovo116915*
C1 C2 C3 C4
0 1 2 3 4
*Stenotrophomonas* denovo124634***
C1 C2 C3 C4
0 10 20 30 40
*Peptoclostridium* denovo68017***
C1 C2 C3 C4
0 1 2 3 4 5
*Clostridium* denovo66475***
C1 C2 C3 C4
0 1 2 3 4
*Delftia* denovo36561***
C1 C2 C3 C4
0 10 20 30 40
*Serratia* denovo124378*
C1 C2 C3 C4
0.00 0.05 0.10 0.15 0.20
*Microbacterium* denovo171745**
C1 C2 C3 C4
0 2 4 6
